# Physiological correlates of a simple saccadic-decision task to extended objects in superior colliculus

**DOI:** 10.1101/2024.03.09.584223

**Authors:** B. Caziot, B. Cooper, M. R. Harwood, R. M. McPeek

## Abstract

Our vision is best only in the center of our gaze, and we use saccadic eye movements to direct gaze to objects and features of interest. We make more than 180,000 saccades per day, and accurate and efficient saccades are crucial for most visuo-motor tasks. Saccades are typically studied using small point stimuli, despite the fact that most real-world visual scenes are composed of extended objects. Recent studies in humans have shown that the initiation latency of saccades is strongly dependent on the size of the target (the “size-latency effect”), perhaps reflecting a tradeoff between the cost of making a saccade to a target and the expected information gain that would result. Here, we investigate the neuronal correlates of the size-latency effect in the macaque superior colliculus. We analyzed the latency variations of saccades to different size targets within a stochastic accumulator model framework. The model predicted a steeper increase in activity for smaller targets compared to larger ones. Surprisingly, the model also predicted an increase in saccade initiation threshold for larger targets. We found that the activity of intermediate-layer SC visuomotor neurons is in close agreement with the model predictions. We also found evidence that these effects may be a consequence of the visual responses of SC neurons to targets of different sizes. These results shed new light on the sources of delay within the saccadic system, a system that we heavily depend upon in the performance of most visuo-motor tasks.

## Introduction

Because vision is most accurate at the fovea, humans constantly move their eyes to refoveate the visual scene, mostly through saccadic eye movements. Although these eye movements are fast and accurate, it has long been observed that they have a surprisingly long initiation latency^1^. This initiation delay is thought to reflect a decision-making process in which an observer accumulates evidence in favor of making an eye-movement. Saccadic eye-movements are costly, both physiologically and functionally^2,3^, thus this saccadic decision-making probably reflects a compromise between visual acuity gained from orienting the fovea to a new location and the cost for moving the fovea. These decisions are among the most common in everyday life, and yet the link between function and decision signal remains poorly understood.

Multiple factors modulating saccade latencies have been well studied, including stimulus contrast^4^, inhibition of return^5^ and the gap effect^6–9^, and the neural bases of these factors have been characterized^10–15^. Target eccentricity is also known to affect latencies, with a minimum for eccentricities between 3-10 deg, steeply rising at shorter eccentricities^16–20^ and more gradually rising at larger eccentricities^19,21^. The neural basis for the dramatic increase in latency for small eccentricity saccade targets has been ascribed either to a fixation zone around the foveal representation in the superior colliculus (SC)^22^, a key structure for saccade generation, or an equilibrium between neurons in the SC^23^.

Unlike arm movements, the kinematics of saccades and their reaction times have typically been thought to be little affected by the size of target objects. Target size has been reported to have little effect on saccade latency or precision^24^, and the scaling of target size to compensate for cortical magnification in the periphery has no effect on saccade latency^25^. Small and inconsistent effects of target size have been reported in other human studies^26,27^. Conversely, a handful of human studies have shown strong effects on saccade latency of the spatial scale of attention ^28,29^ or target size^30,31^, and this phenomenon has been termed the *Size-Latency Effect*^28–31^.

Here, we first establish that the Size-Latency Effect exists in monkeys, as it does in humans, and go on to investigate the involvement of visuomotor neurons in SC in this effect. To do this, we fit a stochastic accumulator decision model to monkeys’ saccade-latency data to infer the values of putative decision-related variables. We then compare the activity of cells in the intermediate layers of the SC to these predicted variables, finding that movement-related activity in the SC closely matches the predictions from the accumulator model. Finally, we show that these effects are most likely caused by the visual responses of the same neurons.

## Results

We trained 2 monkeys (*Macaca mulatta*) to perform saccadic eye movements toward a large ring target. After a cell exhibiting movement-related activity was isolated, we used a delayed-saccade task to map the preferred saccade direction and amplitude associated with the cell, and then displayed ring-shaped targets centered at the cell’s preferred location. Ring sizes were varied from trial-to-trial such that the Distance-to-Size Ratio (DSR=Distance/Size) had one of 4 values ranging between approximately 0.3 and 1.0, with a larger DSR corresponding to a smaller target. Monkeys were rewarded for making a saccade to the center of the rings.

The size of the targets modulated behavior in several ways. Figure 1B plots median saccadic latencies as a function of DSR across sessions for both monkeys. Both monkeys exhibited a significant increase of their saccadic latencies with target size (resampling, see Methods, p<0.001), establishing a strong Size-Latency effect in monkeys (see Figure S1 for latencies plotted as a function of target size). Saccadic latencies ranged from almost 250ms for the largest targets to only about 150ms for the smallest targets. Figure 1C plots mean saccadic error (distance between the eye landing point and the center of the ring target) as a function of DSR. Saccadic errors significantly decreased with target size for both animals (resampling, p<0.001). Finally, Figure 1D plots the mean saccade peak velocity as a function of DSR. Peak velocity significantly decreased with smaller targets for monkey Q (resampling, p<0.001), from approximately 640 to 500deg.sec^-1^, but remained stable at approximately 430deg.sec^-1^ for monkey P (resampling, p=0.47). Therefore, the size-latency effect caused, as in humans^28–31^, a dramatic increase in saccadic latencies for larger targets, as well as larger saccade endpoint spread. In one animal it was also accompanied by a slight decrease in saccade peak velocities but not in the other animal.

**Figure 1:**
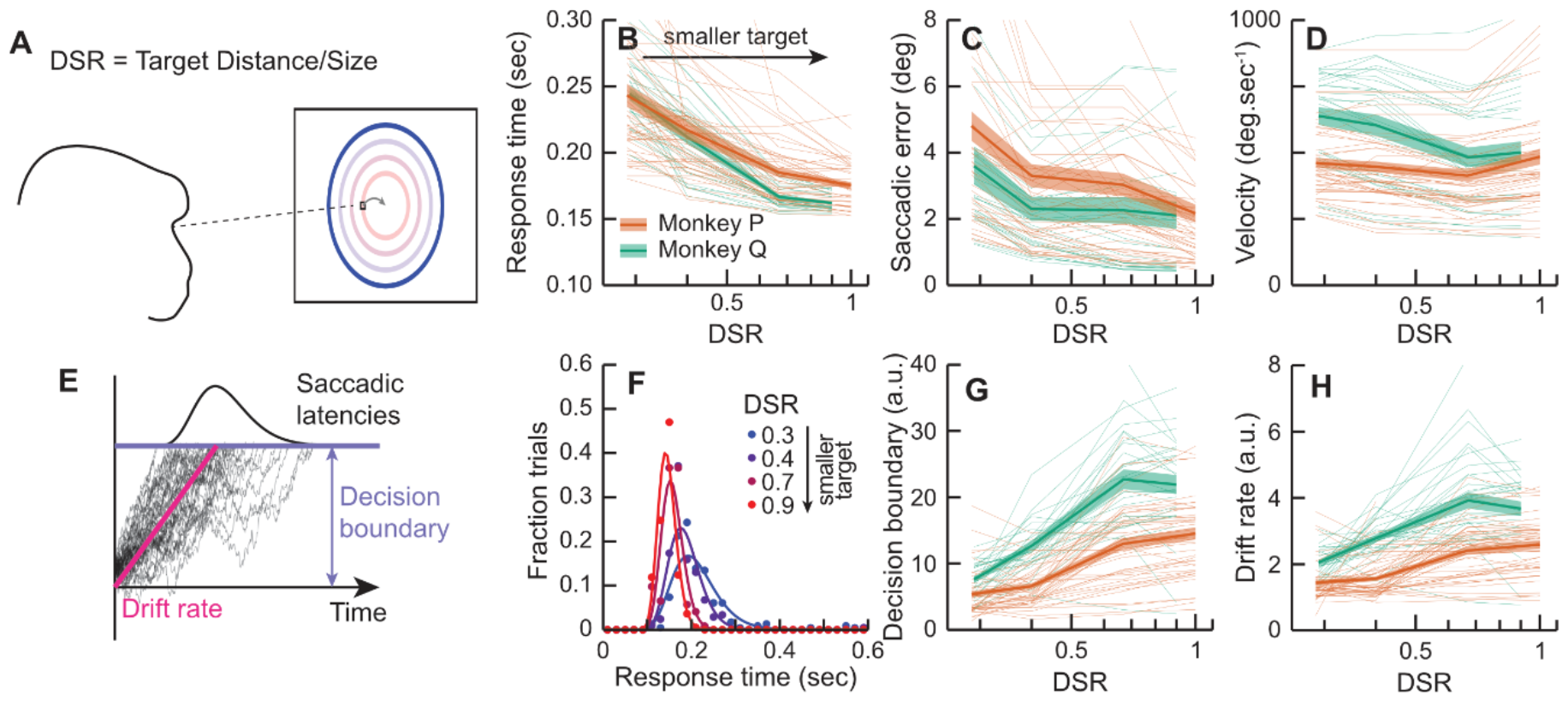
(A) Monkeys were trained to make saccades to targets with different Distance-to-Size Ratios (DSRs). A higher DSR corresponds to a smaller and/or more distant target. Only one target was presented on each trial. (B) Median saccadic latency as a function of DSR for both animals (green and orange). Thin lines correspond to individual sessions, thick lines to mean across sessions and shaded area to standard errors. (C) Mean saccadic error as a function of DSR for both animals. (D) Peak saccadic velocity as a function of DSR for both animals. (E) Saccadic latencies were modeled as the accumulation of a noisy signal with a drift-rate towards a decision boundary. Saccades are triggered when the decision boundary is crossed predicting an inverse-Gaussian distribution of saccadic latencies. (F) Example of distributions of saccadic latencies as a function of DSR (blue to red) for a single session. Dots are fraction of trials within 40ms bins and lines are model fits. (G) Predicted decision boundaries as a function of DSR. (H) Predicted drift-rate as a function of DSR.

We modeled saccadic decisions as an accumulation of a noisy signal towards a decision boundary (see Figure 1E). This class of models is commonly used to model decision time^32^. We assumed the noisy signal to be a Wiener process of mean µ>0 and fixed noise σ=1 (see Figure S2 for a different model leading to similar conclusions). The distribution of crossing times is given by an inverse Gaussian distribution (see Methods). This model corresponds to the continuous limit of Wald’s Sequential Probability Ratio Test against one-sided alternatives^33^, that is where one is preoccupied only with detecting the presence of a target at a fixed α-error rate (false-positives). One-sidedness is warranted here, because the alternative hypothesis would correspond to the animals deciding never to move their eyes.

We fitted inverse Gaussian distributions to the distribution of saccadic latencies using Maximum Likelihood Estimation. Figure 1F shows an example of saccadic latency distributions as a function of DSR (red to blue) for 1 recording session. Lines show that inverse Gaussian distributions fit data well. From the model fit, we can predict the information accumulation rate (drift rate) and height of the decision boundary. Figures 1G,H plot respectively decision boundaries and drift rates as a function of DSR. Predicted drift-rates significantly increase as the target size decreases for both monkeys (resampling, p<0.001), as might be expected from the shorter saccadic latencies associated with smaller targets. The predicted height of the decision boundary also increases significantly as the target size decreases for both monkeys (resampling, p<0.001). This effect is more surprising as an increased decision boundary leads to longer saccadic latencies.

Therefore, modeling saccadic decisions as a stochastic accumulation leads to the (perhaps counter-intuitive) prediction that smaller targets are associated with a higher accumulation rate and increased decision boundary, with a stronger increase in the accumulation rate leading to an overall drop in the saccadic latencies. Note that this modelling (as well as^28^) relies on the assumption of a fixed noise level. Relaxing this assumption leads to an under-specified problem where these effects could be explained by any combination of joint changes in accumulation rate, decisional threshold and noise level.

We then investigated how neuronal activity compares with behavior and predictions from our modeling framework. We first characterized cell activity in a delayed-saccade task (see example cells in Figure S3 and S4). All cells in both monkeys showed a significant increase of activity in the 100ms around saccade-onset as compared to the 100ms prior to target-onset (resampling, see Methods, p<0.05). Most cells (48/51 for Monkey P and 24/28 for Monkey Q) also exhibited a significant increase of activity in the 100ms after target onset compared to 100ms prior to target onset (resampling, p<0.05). As almost all cells exhibited both visual- and saccade-related activity, we computed a Visuo-Motor Index^34^ (see Methods) to place cells on a continuum between pure visual cells (VMI of +1) to pure motor cell (VMI of -1). All cells except one (see Supplementary Material) had a VMI lower than 0, indicating that cell activity in our sample was largely dominated by saccade-related responses.

We then examined cell activity during the Ring task. Because the spread of saccadic endpoints was higher for larger rings than smaller rings, we only analyzed trials where saccade endpoints were within 2 degrees of the target center. Figures 2A and 2C plot the firing rate of 2 example cells as a function of target onset. The cell in figure 2A shows a clear visual response approximately 50ms after target onset for the 2 smaller target sizes (DSRs of 0.7 and 0.9). However, this visual response is almost non-existent for the 2 larger target sizes (DSRs of 0.3 and 0.4). The cell in Figure 2C exhibits a small visual response for the highest DSR and no visual response in other conditions. Figures 2B and 2D show the firing rate of the same 2 cells as a function of time relative to saccade onset. Both cells exhibit a sharp increase in firing rate in the 50ms prior to saccade onset and peak shortly before initiation of the eye-movement, a typical pattern for cells in the Superior Colliculus^35,36^. Both cells show a clearly reduced motor burst for larger targets than smaller targets, even though only trials with similar saccadic endpoints were analyzed. To quantify this effect, for each cell and condition we estimated peak visual activity (maximum firing rate in the 25-75 ms window after target onset), peak motor activity (maximum firing rate in the 50ms preceding saccade onset) and motor build-up rate (increase in rate during the [-40,-10] ms window before saccade onset). These values are plotted in Figures 2F, 2G and 2H respectively. Comparison between the largest and smallest targets reveals an increase from approximately 40 Hz to 100 Hz in peak visual activity, while peak motor activity increased from approximately 70 to 150 Hz. Finally, the change in build-up rate increased from approximately 0.75 to 2 Hz-ms^-1^. To test significance, we resampled our dataset and regressed peaks and build-up rates as a function of DSR for each sample. This analysis confirmed that peak visual activity, peak motor activity and motor build-up rate all increased significantly for both monkeys as a function of DSR (resampling, p<0.001).

**Figure 2:**
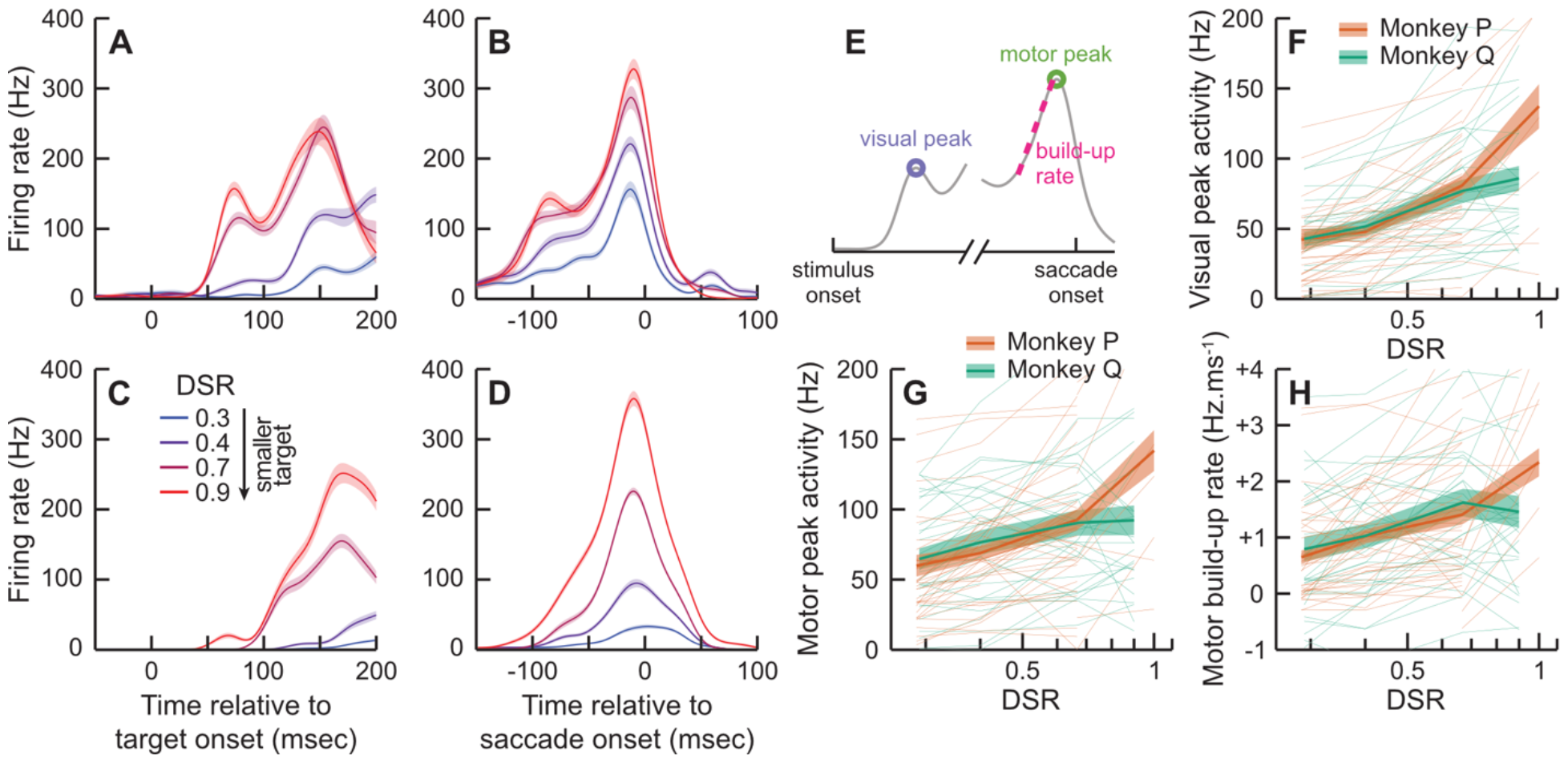
(A) Example of a neuron’s firing rate as a function of time relative to stimulus onset (abscissa) and target DSR (colors). (B) Same as a function of time relative to saccade-onset. (C) and (D) same as (A) and (B) for a different neuron. (E) Depiction of the visual peak, build-up rate and motor peak. (F) Visual peak as a function of DSR (abscissa) for both animals (color). Thin lines are individual sessions. Thick line and shaded areas the mean and SE. (G) Motor peak as a function of DSR. (H) Build-up rate as a function of DSR.

Therefore, as predicted by our model, both peak motor activity and build-up rate increased when the targets became smaller. While this normative approach does not make predictions about visual responses, these strong modulations, comparable in amplitude to the change in motor peak activity, are potentially the source of the size-latency effect at a mechanistic level. To investigate this question, we first looked at the relationship between neural activity and saccadic latencies within condition, that is, for an identical visual stimulus. We binned trials in quintiles of saccadic latency, and computed the same metrics for each latency bin. Figures 3A,B,C plot peak visual activity, peak motor activity and build-up rate, respectively, normalized by the mean firing rate within condition and as a function of the latency bin. For both monkeys, peak visual and motor activity dropped by approximately 20 Hz between the fastest and slowest latency bins. The effect on motor build-up rate is less clear, with a trend towards a small decrease in build-up rate with increasing latency. We tested significance by resampling our dataset and regressing these metrics as a function of latency bins for each sample. Both visual peak activity and motor peak activity significantly decreased as latency increased for both animal (resampling, p<0.001). However, the build-up rate did not significantly decrease for monkey Q (resampling, p=0.16) and just reached significance for monkey P (p=0.047). Consequently, even for identical visual stimuli a higher visual activity was associated with faster saccadic initiation times, and similarly for motor peak activity. But, build-up rate seemed less clearly linked with saccadic initiation time.

**Figure 3:**
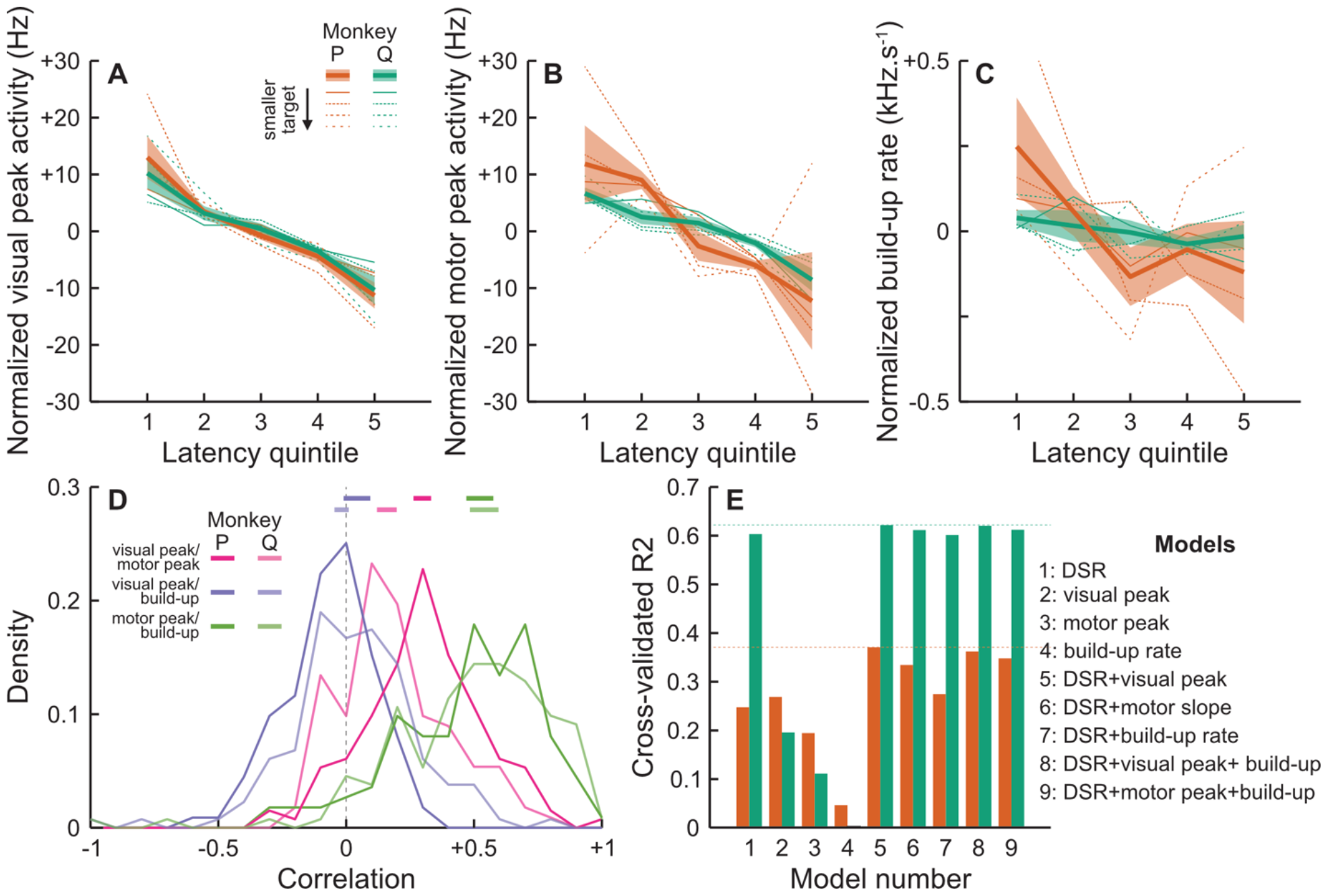
(A) Peak visual activity, normalized by the mean peak activity within condition, as a function of saccadic latency quintiles. (B) Normalized peak motor activity as a function of saccadic latency quintiles. (C) Motor build-up rates as a function saccadic latency quintiles. (D) Distribution of correlation coefficients across trials, within cells and conditions, between visual peak and motor peak activity (pink), between visual peak and build-up rate (purple) and between motor peak and build-up rate (green). Thick horizontal lines at the top indicate 95% confidence interval of the distributions’ median. (E) Cross-validated R^2^ values of various Generalized Linear Models.

Modulations in visual and motor activity at the scale of individual neurons in superior colliculus seem associated with substantial changes in saccadic latencies across conditions (the Size-Latency effect), but also within conditions (for identical visual stimuli). To make this link more explicit, we then investigated the relationship between visual and motor activity from trial to trial. For each trial, we estimated the peak visual activity, peak motor activity and motor build-up rate, and computed their correlations within condition. These correlation factors, plotted in Figure 3D, were consistent across animals. There was a moderate correlation between visual peak and motor peak activity (median correlation 0.31 and 0.16 for monkeys P and Q respectively). The median correlation was significantly higher than 0 for both animals (resampling of the distribution of correlation factors, p<0.0001). The correlation between visual peak activity and motor build-up rate was not significantly different than 0 for both animals (median correlation 0.04 and -0.05 for monkeys P and Q respectively, p=0.90 and 0.98). Finally, the correlation between motor build-up rate and motor peak activity was strongest and significantly different than 0 in both animals (0.54 and 0.53 for monkeys P and Q respectively, p<0.001).

Finally, we tested whether saccadic latencies are better predicted by visual or motor activity. To address this question, we fitted and cross-validated various Generalized Linear Models. The goal of this model comparison was to assess which variables are best predictors of saccadic latencies. Figure 3E shows cross-validated R^2^ values, a measurement of the variance explained by the model. Results are fairly consistent for both monkeys. Unsurprisingly, DSR alone explained a high amount of variance in saccadic latencies. Visual peak activity alone also explained a good amount of variance. These 2 variables together explained a little over 60% of the variance for one animal and almost 40% for the other, better than either variable alone. Remarkably, motor peak activity alone did not explain as much variance as visual peak activity, and the model including both DSR and motor peak activity did not explain more variance than the models including only visual peak activity. Finally, build-up rate alone explained very little variance in saccadic latencies, and including this variable in any model did not improve explained variance.

## Discussion

Here, we investigated the neural underpinning of the Size-Latency effect. Behaviorally, we found, as did prior studies^28–30^, that saccadic latencies were strongly influenced by the Distance-to-Size Ratio. To optimize neural data collection, the animals always performed the same eye-movement to the center of the RF of the cell under study while target size was varied. Yet, the behavioral effect was remarkably similar, in range and amplitude, to the size-latency effect documented in humans. This confirms that the size-latency effect is well-suited for investigating simple saccadic decisions in primates.

We then fitted a stochastic accumulator decision model to the behavioral data. As in our prior study in humans^28^, we predicted a higher accumulation rate and higher decision boundary with smaller targets (higher DSR). These joint effects are counter-intuitive because they go in opposite directions, with a higher accumulation rate leading to faster saccadic latencies, and higher decision boundaries leading to slower saccadic latencies.

We compared these predictions with the actual firing rates of cells in the intermediate layers of SC that exhibit saccade-related activity^22^. As predicted from modelling, we found that SC neurons have a higher build-up rate prior to saccade initiation as well as higher peak motor activity. This result is interesting because the decisional threshold and accumulation rate predicted by the normative stochastic accumulation model are theoretical constructs that can algorithmically be instantiated in different ways ^37^. Here it seems that in the SC these theoretical variables are represented in a near direct fashion by the firing rates of individual neurons. Similar representations in the cortex are often more complex (e.g. ^38^), perhaps reflecting the relative simplicity of SC due to its longer evolutionary history^39^ and relative proximity to the final motor output^40–43^.

However, despite a doubling of the motor peak activity, the velocity of saccadic eye-movements was relatively stable, and even dropped slightly for smaller targets in one animal. This result is surprising because saccadic latencies, spread and velocities have typically been observed to covary with motor activity of SC cells^44–46^.

We also found that visual activity was strongly modulated by the target’s size, with visual activity being almost completely suppressed for the largest ring stimuli. This suppression was to be expected since cells in SC have suppressive surrounds, unlike antagonistic surrounds in other visual areas ^36^. We looked at the relationship between visual and motor activity with saccadic latencies not only across conditions (for different ring sizes), but also within conditions (for identical visual stimuli). We found that even within conditions, higher visual and motor peak activity were associated with shorter saccadic latencies. This result is in contrast with other studies that found a fixed level of activity at the time a motor response is produced both in the cortex^47,48^ and in SC^49–52^. This observation is strengthened, first, by the fact that peak motor activity was correlated with peak visual activity, but not motor build-up rate, and second, from the fact that including build-up rates in a Generalized Linear Model did not improve predictions from the model. The strong modulations observed here, with a doubling of peak motor activity between the smallest and largest DSRs demonstrates that reaching a fixed level of motor activity^51,53,54^ to trigger a saccade is not a general rule in SC, a finding that has also been observed in cortex^55^.

Taken together, these results suggest that the size-latency effect is a direct consequence of the visual responses of cells in SC. Larger targets elicit reduced visual activity over a larger area. This visual activity is followed by a fixed ramp-like increase of firing rate peaking shortly prior saccade onset. Interestingly, the build-up rate was modulated by the target size, but uncorrelated with visual activity within condition. However, the build-up rate was highly correlated with motor peak activity. Cells in SC directly inhibit omnipause neurons^56^, which, in turn, gate the activity of burst neurons in the brainstem. Saccadic velocities are directly related to the magnitude of hyperpolarization of omnipause neurons^57^. We can conclude from the relatively constant saccadic velocities that the overall number of spikes sent to omnipause neurons was also relatively constant across conditions, despite the strong drop in motor peak activity for larger targets. However, we can only speculate about the relationship between build-up rate and DSR. Although the build-up of activity prior to saccade onset is usually modelled as linear for simplicity (e.g. ^47,48,51,52,58^), it actually tends to increase non-linearly, a result of lateral interactions between neurons^47,48^. This would explain why both peak motor activity and build-up rates are reduced for larger targets.

However, our results should be interpreted in the light of possible caveats. First, the distribution of recorded cells was heavily biased towards Visuo-Motor cells. We only selected cells that exhibited saccade-related activity and consequently, we did not observe the visual response of visual-only cells. However, it stands to reason that their visual responses would be comparable.

Second, the visual peak was measured over a fixed 50 ms window during the first 75ms after target onset. Faster saccade latencies are correlated with higher peaks, not shorter periods of visual activity. Although this time window is typically when visual target selection is thought to take place, our data do not by themselves imply that faster latencies are due to faster visual processing. Third, the build-up rate is measured over a smaller 30ms window shortly before the saccade onset, which naturally only captures a snapshot of the build-up accumulation rate.

Visual and motor activity in SC have generally been considered independent^61^. This belief originated from early microstimulation studies which found no effect of stimulation parameters on eye-movements^62,63^, although later studies showed more nuanced results (e.g. ^64^). Similarly, at least in some circumstances visual activity does influence the motor response of individual cells in SC^65,66^. Our results show that another way to make this relationship obvious is to use very large targets. Since these types of elements are omnipresent in real-life visual scenes, rather than small targets on homogenous backgrounds, it is likely that the relationship between visual and motor activity for saccadic eye-movements generation has been under-appreciated in prior studies.

Finally, we should note that if the Size-Latency effect is caused by the visual response of cells in SC, as suggested by our results, this effect cannot be fully accounted for by the SC alone. Indeed, studies in humans^28,29^ used stimuli consisting of a small ring target inside a larger ring target, and observers were instructed to pay attention to the small or large target. This stimulus should elicit the same visual response in SC. Yet our results predict that the visual response would instead depend on which part of the stimulus observers pay attention to, suggesting that this task may be well suited for investigating attentional gating of visual signals by the cortex to SC.

## Methods

### Physiological apparatus

All experimental protocols were approved by the Institutional Animal Care and Use Committee at the State University of New York College of Optometry and complied with the guidelines of the Public Health Service *Policy on Humane Care and Use of Laboratory Animals*. A head-holder system and stainless-steel recording chamber to access the SC bilaterally were implanted under isoflurane anesthesia and aseptic surgical conditions in two rhesus monkeys (*Macaca mulatta*). Antibiotics (cefazolin sodium) and analgesics (buprenorphine hydrochloride) were administered as needed during the recovery period under the direction of a veterinarian.

In each recording session, a tungsten microelectrode (FHC, Inc.) with impedance ranging from 1 to 2 MΩ at 1 kHz was lowered into SC using a motorized microdrive (NAN Instruments, Ltd.). A Plexon Multichannel Acquisition Processor system (Plexon, Inc.) amplified and band-pass filtered the microelectrode signal and was used to identify action potentials. Spike-sorting was verified offline using the Plexon Offline Sorter analysis software. Only well-isolated single units were included in the analyses.

### Behavioral apparatus

Testing was performed in a dimly illuminated room. Experimental control, data acquisition, and presentation of visual displays were carried out by a custom real-time MATLAB program on a Macintosh G4 computer using the Psychophysics Toolbox^67–69^. Visual stimuli were presented on a monitor positioned 29 cm in front of the monkeys. The monitor had a spatial resolution of 800 x 600 pixels and a refresh rate of 75 Hz. Eye position was sampled at 1 kHz using an EyeLink 1000 infrared video tracker (SR Research).

### Procedure

In experimental sessions, a microelectrode was lowered using a stainless-steel guide-tube to locations in the SC which had been previously mapped. Once a cell was well isolated, its preferred saccade direction and amplitude were estimated in a delayed-saccade task by manually varying the location of the target across trials. Only cells showing saccade-related activity in the delayed-saccade task were recorded in the subsequent Ring Task. The average number of trials collected per session for the Ring Task was 400 for monkey P and 396 for monkey Q.

### Delayed-saccade task

A white square fixation point subtending 0.25° with a luminance of 1.5 cd/m^2^ appeared in the central position against a homogenous dim background of 0.2 cd/m^2^. Monkeys were required to keep their eyes within a 1–1.5° window around the fixation point during an initial fixation interval of 450–650 ms. At the end of this interval, the fixation point remained illuminated and a target stimulus was presented at a peripheral location aligned with the RF of the cell. Monkeys were required to maintain fixation until the disappearance of the fixation point approximately 500 ms later. Once the fixation point disappeared, the monkeys were rewarded for making a saccade to the peripheral stimulus. Eye-position tolerance windows around the target were equal to the stimulus eccentricity divided by 5.

### Ring Task

Once we located the center of a cell’s movement field, we presented ring stimuli centered at the cell’s preferred location. We varied the size of the rings to produce varying values of Distance-to-Size Ratio (DSR). The rings had a luminance of 12.4 cd/m^2^ and a thickness equal to the eccentricity of the ring center divided by 20, and were presented on background of 0.2 cd/m^2^. Monkeys were rewarded for making a saccade to the center of the ring. The eye-position tolerance window around the ring center was equal to the eccentricity of the ring center divided by 5. For monkey Q DSR values were always 0.29, 0.40, 0.67 and 0.90. For monkey P they were 0.29, 0.40, 0.67 and □1. Because the Distance-to-Size latency effect saturates at a DSR value of 1^28^, we pooled all DSR values higher than 1 and plot them as 1 in Figures for clarity.

### Cell classification

We characterized the relative strength of the visual and movement-related responses of the cells with a Visuo-Motor Index (VMI^34^) calculated using data from the delayed-saccade task. First, we estimated whether each cell had saccade-related activity by comparing activity in the 100ms prior to saccade initiation to 100ms prior to target onset. Second, we estimated whether cells had a visual response by comparing activity in the 100ms range after target onset to the 100ms prior to target onset. Significance was established by resampling 1,000,000 times the difference in mean firing-rate between 2 event windows. VMI was then computed as:

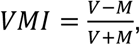

where V and M are the peak visual (V) and motor (M) activity in the aforementioned windows. This index varies from -1 to +1, with -1 indicating a cell that has only have a motor response, and +1 a cell that has only have a visual response, and 0 a cell with visual and motor responses that are equal in magnitude. The goal of this index is to capture the continuum between so-called visual neurons, motor neurons and visuo-motor neurons.

### Data Analysis

Offline data analysis was performed using custom MATLAB (Mathworks) scripts. Saccades were detected using velocity and acceleration criteria. To generate continuous spike-density functions, neural events were convolved with an acausal Gaussian filter (σ=5ms). For statistical reliability, only conditions having at least 10 trials were included in the analyses (84% and 100% of conditions for monkeys P and Q respectively). When assessing statistical significance, we adopted a criterion α-level of 0.05. Unless specified otherwise, statistical testing was done by resampling, either individual trials within a session or the mean across sessions, 1,000,000 times, and re-estimating the tested parameter for each sample (e.g. the slope parameter for a linear regression). Unless otherwise specified, all tests were two-sided.

For model comparison, for each cell separately we randomly selected 20% of the dataset as a validation set. We fitted a GLM to the remaining 80% of the data, then computed the adjusted R^2^ values of the model on the validation set, according to:

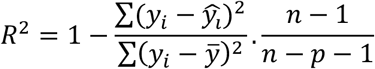

With *n* and *p* the number of observations and number of model parameters respectively. We performed this comparison 1,000,000 times and computed the mean R^2^ value across cells and samples. We regressed inverse saccadic latencies with DSR values as categorical variables (4 groups), peak visual activity, peak motor activity and motor build-up rate.

### Accumulator model

We modeled decisions as the accumulation of a noisy signal toward a decision boundary. A saccade is initiated when the decision boundary is crossed. We assumed that the accumulated signal is generated by a Wiener process with drift *μ* and variance *σ*^2^

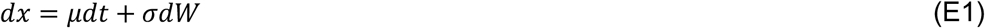

Therefore, the probability for the accumulator to reach the decision bound *z* at a time *t* follows an inverse Gaussian distribution^70^:

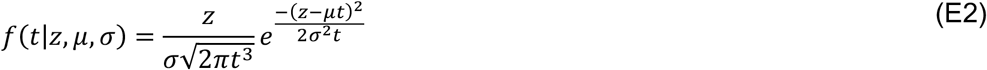

We used Maximum Likelihood Estimation to fit the distribution of saccadic latencies with the predicted distribution:

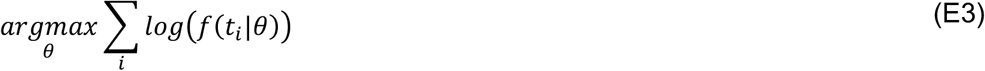

For comparison, we also fitted a LATER saccadic decision model^58^. Results across models were highly correlated (see Figure S2) and all results described here remain valid using either model.

## Acknowledgments

This research was supported by a National Science Foundation grant (1232654) to M.R. Harwood, a National Institutes of Health grant (R01-EY030669) to R.M. McPeek, and a Deutsche Forschungsgemeinschaft (DFG, German Research Foundation) grant (524696675) to B. Caziot. We thank Junghyun Park, PhD for expert assistance with training and data collection.

## References

1. Carpenter, R.H.S. (1981). Eye movements: Cognition and visual perception. Oculomot. Procrastination Fish. DF Monty RA Send. JW Eds Pp 237, 246.

2. Harris, C.M., and Wolpert, D.M. (2006). The Main Sequence of Saccades Optimizes Speed-accuracy Trade-off. Biol. Cybern. 95, 21–29. 10.1007/s00422-006-0064-x.

3. Matin, E. (1974). Saccadic suppression: a review and an analysis. Psychol. Bull. 81, 899.

4. Ludwig, C.J., Gilchrist, I.D., and McSorley, E. (2004). The influence of spatial frequency and contrast on saccade latencies. Vision Res. 44, 2597–2604.

5. Klein, R.M. (2000). Inhibition of return. Trends Cogn. Sci. 4, 138–147.

6. Saslow, M.G. (1967). Effects of components of displacement-step stimuli upon latency for saccadic eye movement. Josa 57, 1024–1029.

7. Reuter-Lorenz, P.A., Hughes, H.C., and Fendrich, R. (1991). The reduction of saccadic latency by prior offset of the fixation point: An analysis of the gap effect. Percept. Psychophys. 49, 167–175. 10.3758/BF03205036.

8. Weber, H., Aiple, F., Fischer, B., and Latanov, A. (1992). Dead zone for express saccades. Exp. Brain Res. 89. 10.1007/BF00229018.

9. McPeek, R.M., and Schiller, P.H. (1994). The effects of visual scene composition on the latency of saccadic eye movements of the rhesus monkey. Vision Res. 34, 2293–2305. 10.1016/0042-6989(94)90108-2.

10. Maunsell, J.H., and Gibson, J.R. (1992). Visual response latencies in striate cortex of the macaque monkey. J. Neurophysiol. 68, 1332–1344. 10.1152/jn.1992.68.4.1332.

11. Lennie, P. (1981). The physiological basis of variations in visual latency. Vision Res. 21, 815–824.

12. Dorris, M.C., Klein, R.M., Everling, S., and Munoz, D.P. (2002). Contribution of the primate superior colliculus to inhibition of return. J. Cogn. Neurosci. 14, 1256–1263. 10.1162/089892902760807249.

13. Dorris, M.C., and Munoz, D.P. (1995). A neural correlate for the gap effect on saccadic reaction times in monkey. J. Neurophysiol. 73, 2558–2562.

14. Mirpour, K., Arcizet, F., Ong, W.S., and Bisley, J.W. (2009). Been there, seen that: a neural mechanism for performing efficient visual search. J. Neurophysiol. 102, 3481–3491. 10.1152/jn.00688.2009.

15. Conroy, C., Nanjappa, R., and McPeek, R.M. (2023). Inhibitory tagging in the superior colliculus during visual search. J. Neurophysiol. 130, 824–837. 10.1152/jn.00095.2023.

16. Wyman, D., and Steinman, R.M. (1973). Latency characteristics of small saccades. Vision Res.

17. Kalesnykas, R.P., and Hallett, P.E. (1994). Retinal eccentricity and the latency of eye saccades. Vision Res. 34, 517–531.

18. Kalesnykas, R.P., and Hallett, P.E. (1996). Fixation conditions, the foveola and saccadic latency. Vision Res. 36, 3195–3203.

19. Hafed, Z.M., and Goffart, L. (2020). Gaze direction as equilibrium: more evidence from spatial and temporal aspects of small-saccade triggering in the rhesus macaque monkey. J. Neurophysiol. 123, 308–322. 10.1152/jn.00588.2019.

20. Poletti, M., Intoy, J., and Rucci, M. (2020). Accuracy and precision of small saccades. Sci. Rep. 10, 16097.

21. Becker, W. (1989). Metrics. In The Neurobiology of Saccadic Eye Movements (Elsevier), pp. 13–67.

22. Munoz, D.P., and Wurtz, R.H. (1995). Saccade-related activity in monkey superior colliculus. I. Characteristics of burst and buildup cells. J. Neurophysiol. 73, 2313–2333.

23. Hafed, Z.M., Goffart, L., and Krauzlis, R.J. (2009). A Neural Mechanism for Microsaccade Generation in the Primate Superior Colliculus. Science 323, 940–943. 10.1126/science.1166112.

24. Kowler, E., and Blaser, E. (1995). The accuracy and precision of saccades to small and large targets. Vision Res. 35, 1741–1754.

25. Yates, D.J., and Stafford, T. (2011). Insights into the Function and Mechanism of Saccadic Decision Making From Targets Scaled By an Estimate of the Cortical Magnification Factor. Cogn. Comput. 3, 89–93. 10.1007/s12559-010-9075-y.

26. Ploner, C.J., Ostendorf, F., and Dick, S. (2004). Target size modulates saccadic eye movements in humans. Behav. Neurosci. 118, 237.

27. Dick, S., Ostendorf, F., Kraft, A., and Ploner, C.J. (2004). Saccades to spatially extended targets: the role of eccentricity. Neuroreport 15, 453–456.

28. Harwood, M.R., Madelain, L., Krauzlis, R.J., and Wallman, J. (2008). The Spatial Scale of Attention Strongly Modulates Saccade Latencies. J. Neurophysiol. 99, 1743–1757.

29. Madelain, L., Krauzlis, R.J., and Wallman, J. (2005). Spatial deployment of attention influences both saccadic and pursuit tracking. Vision Res. 45, 2685–2703.

30. De Vries, J.P., Azadi, R., and Harwood, M.R. (2016). The saccadic size-latency phenomenon explored: Proximal target size is a determining factor in the saccade latency. Vision Res. 129, 87–97.

31. Vullings, C., Harwood, M.R., and Madelain, L. (2019). Reinforcement reduces the size– latency phenomenon: A cost–benefit evaluation of saccade triggering. J. Vis. 19, 16–16.

32. Luce, R.D. (1986). Response times: Their role in inferring elementary mental organization (Oxford University Press on Demand).

33. Wald, A. (2004). Sequential analysis (Courier Corporation).

34. Massot, C., Jagadisan, U.K., and Gandhi, N.J. (2019). Sensorimotor transformation elicits systematic patterns of activity along the dorsoventral extent of the superior colliculus in the macaque monkey. Commun. Biol. 2, 1–14. 10.1038/s42003-019-0527-y.

35. Gandhi, N.J., and Katnani, H.A. (2011). Motor Functions of the Superior Colliculus. Annu. Rev. Neurosci. 34, 205–231. 10.1146/annurev-neuro-061010-113728.

36. Wurtz, R.H., and Albano, J.E. (1980). Visual-motor function of the primate superior colliculus. Annu. Rev. Neurosci. 3, 189–226.

37. Marr, D. (2010). Vision: A computational investigation into the human representation and processing of visual information (MIT press).

38. Ditterich, J. (2006). Stochastic models of decisions about motion direction: behavior and physiology. Neural Netw. 19, 981–1012.

39. Cooper, B., and McPeek, R.M. (2021). Role of the superior colliculus in guiding movements not made by the eyes. Annu. Rev. Vis. Sci. 7, 279–300.

40. Sparks, D.L., and Hartwich-Young, R. (1989). The deep layers of the superior colliculus. Rev. Oculomot. Res. 3, 213–255.

41. Keller, E.L., McPeek, R.M., and Salz, T. (2000). Evidence against direct connections to PPRF EBNs from SC in the monkey. J. Neurophysiol. 84, 1303–1313.

42. Rodgers, C.K., Munoz, D.P., Scott, S.H., and Paré, M. (2006). Discharge properties of monkey tectoreticular neurons. J. Neurophysiol. 95, 3502–3511. 10.1152/jn.00908.2005.

43. Miyashita, N., and Hikosaka, O. (1996). Minimal synaptic delay in the saccadic output pathway of the superior colliculus studied in awake monkey. Exp. Brain Res. 112, 187–196. 10.1007/BF00227637.

44. Dorris, M.C., and Munoz, D.P. (1998). Saccadic probability influences motor preparation signals and time to saccadic initiation. J. Neurosci. 18, 7015–7026.

45. Sparks, D.L., Holland, R., and Guthrie, B.L. (1976). Size and distribution of movement fields in the monkey superior colliculus. Brain Res. 113, 21–34. 10.1016/0006-8993(76)90003-2.

46. Wurtz, R.H., and Goldberg, M.E. (1972). Activity of superior colliculus in behaving monkey. 3. Cells discharging before eye movements. J. Neurophysiol. 35, 575–586. 10.1152/jn.1972.35.4.575.

47. Roitman, J.D., and Shadlen, M.N. (2002). Response of neurons in the lateral intraparietal area during a combined visual discrimination reaction time task. J. Neurosci. 22, 9475–9489.

48. Hanes, D.P., and Schall, J.D. (1996). Neural control of voluntary movement initiation. Science 274, 427–430.

49. Horwitz, G.D., and Newsome, W.T. (2001). Target Selection for Saccadic Eye Movements: Prelude Activity in the Superior Colliculus During a Direction-Discrimination Task. J. Neurophysiol. 86, 2543–2558. 10.1152/jn.2001.86.5.2543.

50. Horwitz, G.D., and Newsome, W.T. (1999). Separate Signals for Target Selection and Movement Specification in the Superior Colliculus. Science 284, 1158–1161. 10.1126/science.284.5417.1158.

51. Ratcliff, R., Cherian, A., and Segraves, M. (2003). A Comparison of Macaque Behavior and Superior Colliculus Neuronal Activity to Predictions From Models of Two-Choice Decisions. J. Neurophysiol. 90, 1392–1407. 10.1152/jn.01049.2002.

52. Jun, E.J., Bautista, A.R., Nunez, M.D., Allen, D.C., Tak, J.H., Alvarez, E., and Basso, M.A. (2021). Causal role for the primate superior colliculus in the computation of evidence for perceptual decisions. Nat. Neurosci. 24, 1121–1131. 10.1038/s41593-021-00878-6.

53. Kim, B., and Basso, M.A. (2008). Saccade target selection in the superior colliculus: a signal detection theory approach. J. Neurosci. 28, 2991–3007.

54. Ratcliff, R., Hasegawa, Y.T., Hasegawa, R.P., Smith, P.L., and Segraves, M.A. (2007). Dual Diffusion Model for Single-Cell Recording Data From the Superior Colliculus in a Brightness-Discrimination Task. J. Neurophysiol. 97, 1756–1774. 10.1152/jn.00393.2006.

55. Heitz, R.P., and Schall, J.D. (2012). Neural mechanisms of speed-accuracy tradeoff. Neuron 76, 616–628. 10.1016/j.neuron.2012.08.030.

56. Raybourn, M.S., and Keller, E.L. (1977). Colliculoreticular organization in primate oculomotor system. J. Neurophysiol. 40, 861–878.

57. Yoshida, K., Iwamoto, Y., Chimoto, S., and Shimazu, H. (1999). Saccade-Related Inhibitory Input to Pontine Omnipause Neurons: An Intracellular Study in Alert Cats. J. Neurophysiol. 82, 1198–1208. 10.1152/jn.1999.82.3.1198.

58. Noorani, I., and Carpenter, R.H.S. (2016). The LATER model of reaction time and decision. Neurosci. Biobehav. Rev. 64, 229–251. 10.1016/j.neubiorev.2016.02.018.

59. Amari, S. (1977). Dynamics of pattern formation in lateral-inhibition type neural fields. Biol. Cybern. 27, 77–87. 10.1007/BF00337259.

60. Trappenberg, T.P., Dorris, M.C., Munoz, D.P., and Klein, R.M. (2001). A model of saccade initiation based on the competitive integration of exogenous and endogenous signals in the superior colliculus. J. Cogn. Neurosci. 13, 256–271.

61. Goldberg, M.E., Eggers, H.M., and Gouras, P. (1991). The oculomotor system. Princ. Neural Sci., 660–676.

62. Robinson, D.A. (1972). Eye movements evoked by collicular stimulation in the alert monkey. Vision Res. 12, 1795–1808.

63. Schiller, P.H., and Stryker, M. (1972). Single-unit recording and stimulation in superior colliculus of the alert rhesus monkey. J. Neurophysiol. 35, 915–924.

64. Stanford, T.R., Freedman, E.G., and Sparks, D.L. (1996). Site and parameters of microstimulation: evidence for independent effects on the properties of saccades evoked from the primate superior colliculus. J. Neurophysiol. 76, 3360–3381. 10.1152/jn.1996.76.5.3360.

65. Edelman, J.A., and Goldberg, M.E. (2001). Dependence of saccade-related activity in the primate superior colliculus on visual target presence. J. Neurophysiol. 86, 676–691. 10.1152/jn.2001.86.2.676.

66. Edelman, J.A., and Goldberg, M.E. (2003). Saccade-related activity in the primate superior colliculus depends on the presence of local landmarks at the saccade endpoint. J. Neurophysiol. 90, 1728–1736. 10.1152/jn.00016.2003.

67. Brainard, D.H., and Vision, S. (1997). The psychophysics toolbox. Spat. Vis. 10, 433–436.

68. Pelli, D.G., and Vision, S. (1997). The VideoToolbox software for visual psychophysics: Transforming numbers into movies. Spat. Vis. 10, 437–442.

69. Kleiner, M., Brainard, D., and Pelli, D. (2007). What’s new in Psychtoolbox-3. Perception 36, 1.

70. Redner, S. (2001). A guide to first-passage processes (Cambridge university press).

