## Supplementary Figures S1-S4 for "Physiological correlates of a simple saccadic-decision task to extended objects in superior colliculus"

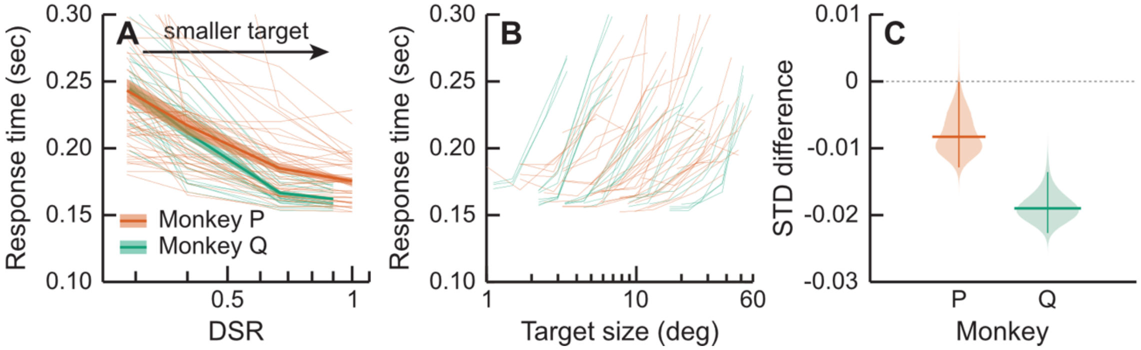


**Figure S1:** (A) Same as Figure 1B: median saccadic latency as a function of DSR for both animals (green and orange). Thin lines are individual sessions, thick lines mean across sessions and shaded area standard error. (B) Same as A but plotted as a function of target size (abscissa). (C) Difference between the standard deviation within 4 bins when saccadic latencies are grouped by DSR as compared to when they are grouped by Target Size.

Evidence that saccadic latencies are better explained by the target Distance to Size Ratio (DSR), rather than just Target Size. Figure S1A plots saccadic latencies as a function of DSR, and Figure S1B as a function of Target Size. Each line corresponds to a different recording session where target eccentricity (and therefore saccadic amplitude) was fixed. DSR was consequently directly related to Target Size. To estimate if saccadic latencies were better explained by DSR than target size, we binned median latencies in 4 bins as a function of either DSR or Target Size and computed the mean standard deviation of median latencies across bins. For both animals the standard deviation of median latencies was smaller when binned by DSR than by Target Size.

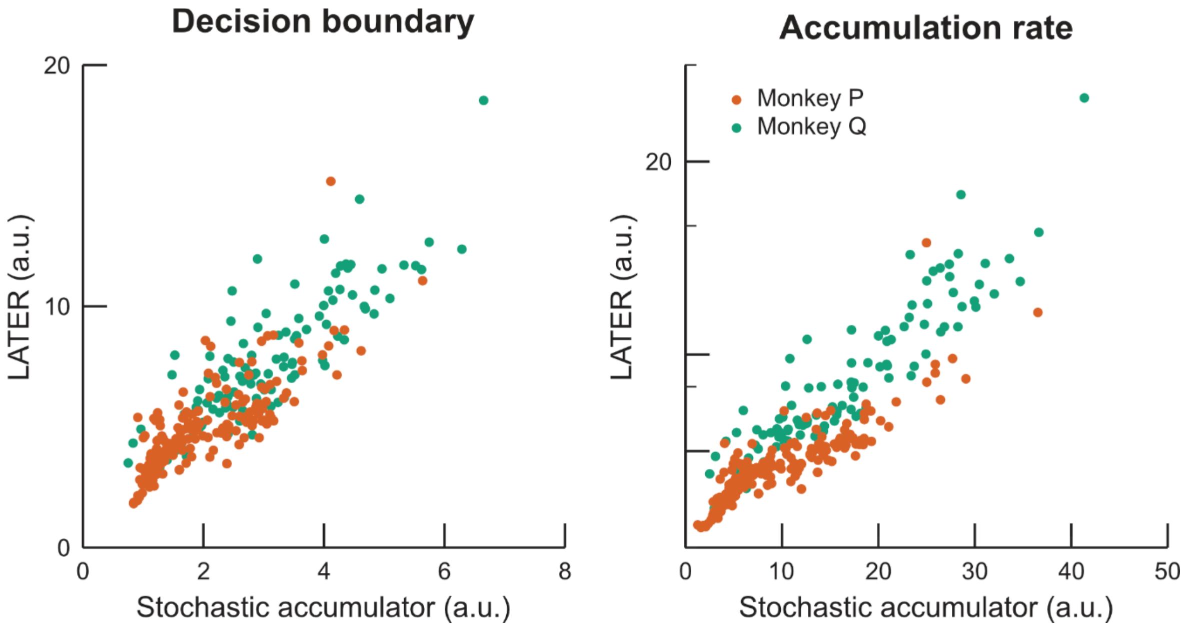


**Figure S2:** Left: Decision boundaries predicted from fitting the LATER model as a function of decision boundaries predicted from the stochastic accumulator model for both animals (colors). Right: Same for accumulation rates.

For each session we fitted a stochastic accumulator model and a LATER model [28,58]. Both models assume the accumulation of a signal towards a decision boundary. The LATER model predicts a reci-normal distribution and the stochastic accumulator predicts an inverse-Gaussian distribution, which are very similar. Indeed, parameters predicted from either model were highly correlated: 0.91 and 0.92 for the accumulation rate for monkeys P and Q respectively; 0.81 and 0.85 for the decision boundary. All results described in the main text remain valid using either model.

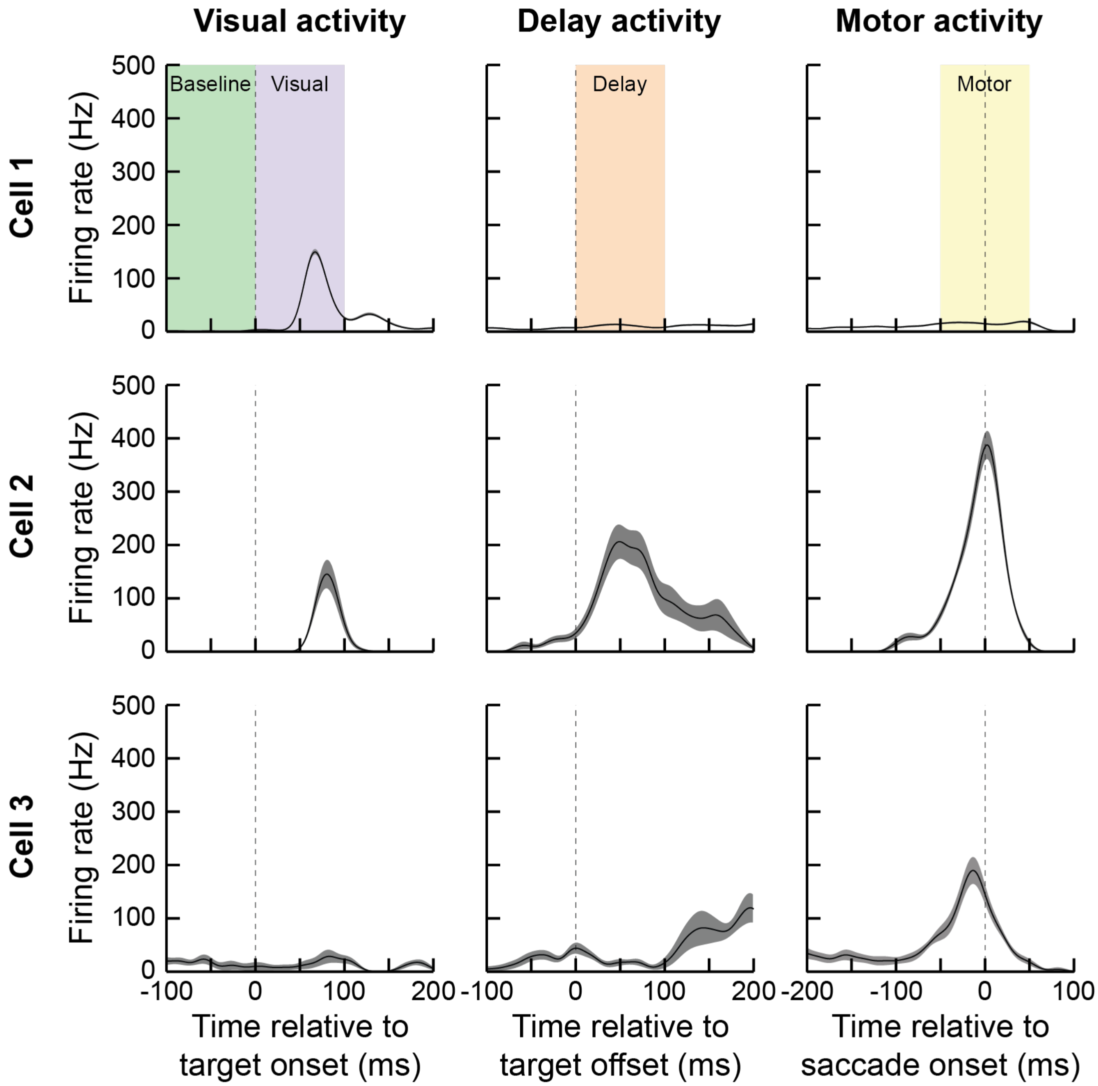


**Figure S3:** Example responses in the delayed saccade task for 3 neurons (rows). The left columns plots mean firing rate (black line) and standard error (shaded area) as a function of time relative to target onset. Middle column relative to target offset and right column relative to saccade onset.

We classified cells with a delayed-saccade task. This task allows dissociating visual from motor activity. Figure S3 shows 3 example cells. The first cell (top row) exhibits a clear increase in firing rate between 50 and 100ms after target onset (left graph), but no change in activity when the target disappears (middle) or when the animal performs the eye-movement (right). The second cell (middle row) also exhibits a clear visual response, as well as an increase in firing rate when the target disappears, and a strong increase in firing rate peaking approximately at eye-movement onset. The last row shows a cell that has only motor activity.

To test whether a cell exhibited visual, delay or motor activity, we computed the mean firing rate within 3 time windows (shaded color areas in Figure S3) and compared this activity to the baseline activity of the neuron in the 100ms prior target onset.

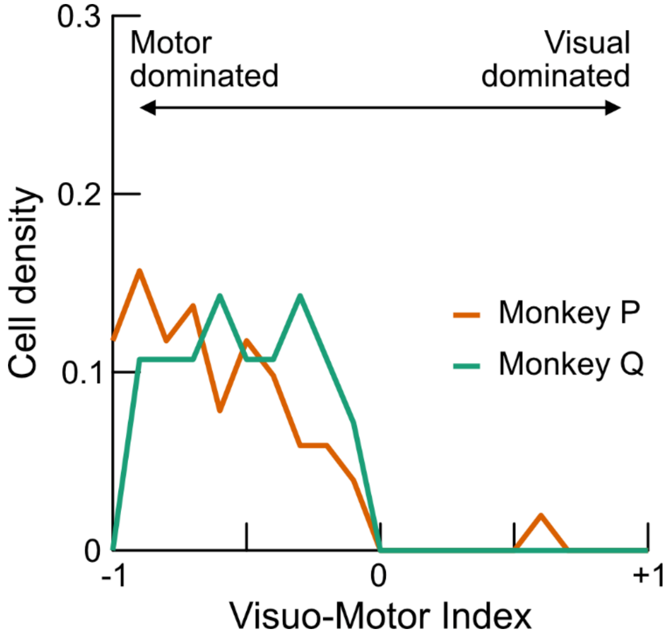


**Figure S4:** Histogram of Visual-Motor Indices for all cells of both animals (color). A value of -1 indicates a pure motor cell (clear motor response, no visual response) and a value of +1 indicates a pure visual cell (clear visual response, no motor response).
